# Development of a Novel Hybprinter-SAM for Functionally Graded Tissue Engineering Constructs with Patterned and Localized Biochemical Signals

**DOI:** 10.1101/2024.11.27.620700

**Authors:** Jiannan Li, Carolyn Kim, Hossein V. Alizadeh, Shreya Garg, Arnaud Bruyas, Peng Zhao, Isadora S. D. Passos, Andrea S. Flores Pérez, Yunzhi P. Yang

## Abstract

Engineering native-mimetic tissue constructs is challenging due to their intricate biological and structural gradients. To address this, Hybprinter-SAM was developed by integrating three bioprinting technologies: syringe extrusion (SE), acoustic droplet ejection (ADE) and molten material extrusion (MME). This system enables not only the creation of mechanical gradients by combining soft and rigid materials but also precise patterning and controlled localization of biochemical signals within printed scaffolds. This capability is beneficial in replicating the complexity of native tissues to enhance functionality. Both the printing process and biomaterials were optimized to balance printability, mechanical integrity, and biocompatibility. As a proof of concept, Hybprinter-SAM was used in a bone-tendon regeneration study to engineer a multi-material construct with patterned fibroblast growth factor 2 (FGF-2), resulting in markers indicative of fibrocartilage development. These findings highlight the potential of Hybprinter-SAM as a versatile platform for diverse tissue engineering applications that require complex, functionally graded tissue constructs.

## INTRODUCTION

Tissue engineering is an interdisciplinary field that combines principles from biology, engineering, and materials science to create biological tissues that can replace, repair, or enhance the function of damaged or diseased tissues in the body ^1, 2^. The goal of tissue engineering is to develop functional tissue constructs that can restore normal function, help in healing, or even create organs for transplantation. This typically involves using a combination of cells, scaffolding biomaterials, and biologically active molecules to guide the growth and remodeling of new tissues ^3^. 3D bioprinting has long been a tool in tissue engineering with its ability to fabricate complex and anatomically relevant constructs. Significant progress has been achieved in engineering tissue constructs with advanced 3D bioprinting techniques and corresponding biomaterials, including molten material extrusion (MME) for thermoplastics ^4–6^, syringe extrusion (SE) for extrudable hydrogels ^7^, stereolithography (SLA) for photocrosslinkable materials ^8, 9^, or droplet printing for bioreagents ^10–12^. Despite these discoveries, engineering functionally graded complex tissues remains challenging, as each printing technique is typically limited to one type of material. Tissues, or multi-tissue interfaces, are often complex constructs comprising of mechanical, compositional, and biological gradients. For instance, the mechanical property of different tissues within the human body spans as much as 7 orders of magnitude ^13^, while the mechanical properties of each family of biomaterials compatible with a specific bioprinting mechanism is often limited (**Figure 1a**). Therefore, there is an inherent demand for advanced engineering tools to fabricate such functionally graded complex tissues.

**Figure 1.**
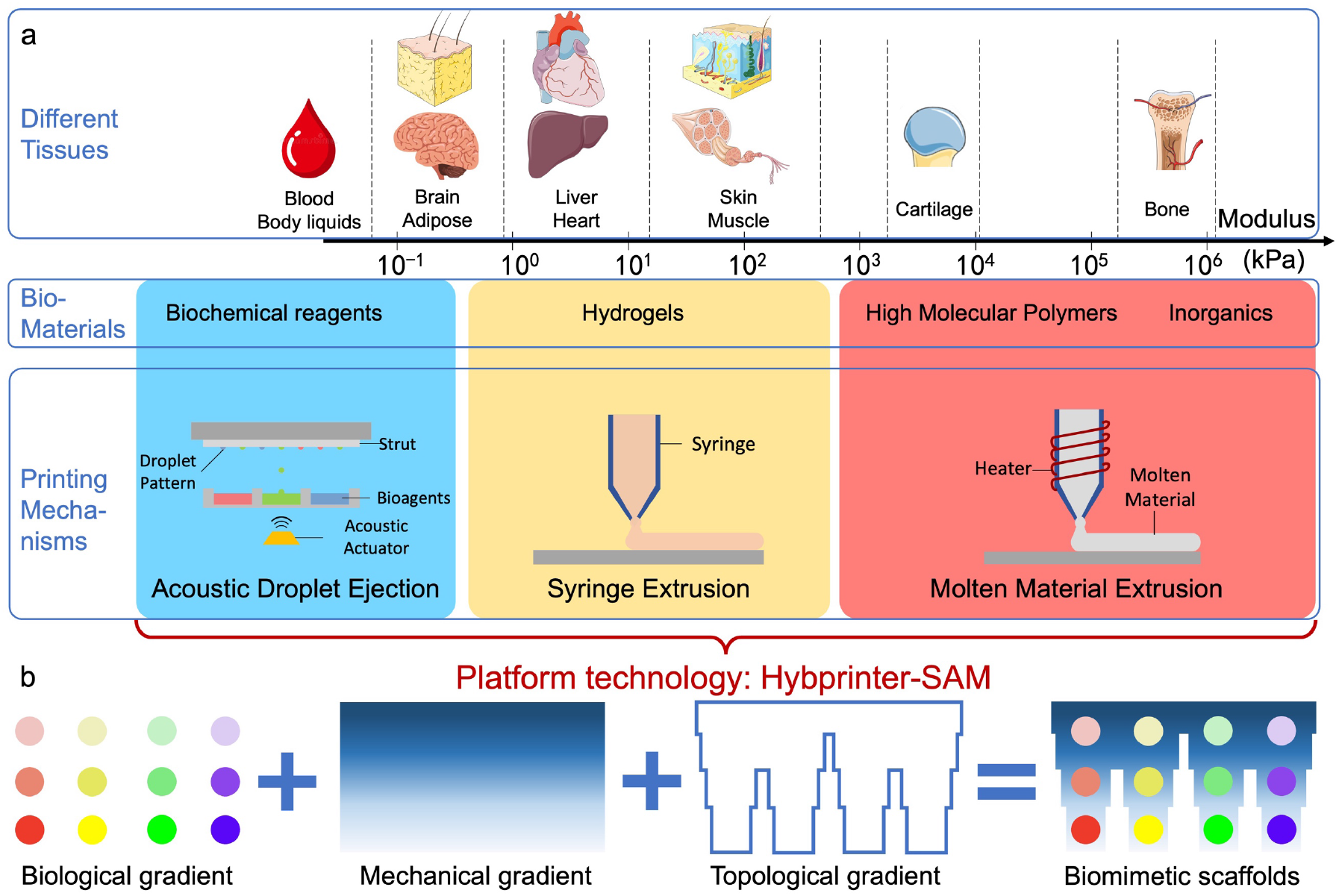
(a) Background and rationale of Hybrid bioprinting in engineering complex tissue interfaces. (b) Premise of Hybprinter-SAM comprising of biological, mechanical and topographical gradient.

One avenue for fabricating such graded complex tissues that researchers have been pursuing is hybrid bioprinting (hybprinting), which employs multiple printing techniques to support a variety of biomaterials. Researchers have been developing hybprinters since the early 2010’s, with an initial common strategy of combining MME and SE for extrusion of rigid and soft biomaterials, respectively, pioneered by D. Cho and J. Malda ^14, 15^. Although similar hybrid designs can be recreated by simply casting a photocrosslinkable hydrogel over a polymer scaffold ^16–18^, the potential of 3D bioprinting reaches far more complex topographical designs. For instance, A. Atala and colleagues built a platform integrating MME and SE modules that could engineer biomimetic human-scale tissues by introducing a sacrificial material to support more complicated patterns ^19^. Other than the combination of MME and SE, additional bioprinting mechanisms have also been introduced to hybprinting, such as droplet printing and SLA. With regards to hybprinting with droplet printing, A. Atala’s group has investigated layer-by-layer patterns by inkjet printing a cell-laden hydrogel layer followed by an electrospun polymer layer ^20^; D. Kelly’s group has also utilized inkjet printing for cell deposition in combination with melt electrowriting for super fine structural support ^21^; and S. Jung et al. used SE to print collagen and dermal fibroblasts and inkjet printing for epidermal keratinocytes to fabricate skin models ^22^. In these hybprinting platforms, droplet printing was predominantly used for cell deposition due to its ability to precisely pattern solutions. Such capability can also be used to deposit other solutions including biomolecules ^23–25^ or hydrogel bioinks ^20, 26, 27^. Hybprinting with SLA mainly aims to utilize its advantage of high-speed and high-resolution printing to create hydrogel constructs with complex geometries. H. Qi and co-workers have developed a hybrid additive manufacturing platform by integrating modified SE and digital light processing-stereolithography (DLP-SLA) for various potential biomedical applications ^28, 29^. By combining DLP-SLA with microfluidic techniques for changing materials, Y. Zhang et al. has achieved functionally graded multi-material printing with DLP-SLA alone ^30^. As these groups continue to explore hybprinting technologies for various applications with a particular interest in musculoskeletal tissue engineering ^31–38^, others have joined the effort in developing integrated 3D printing systems for complex tissue engineering ^39, 40^. Commercialized hybprinters have also been developed with improved robustness and scaling capability ^41–43^. The efforts made in both academia and industry highlight the importance of hybprinting. Among these hybprinting techniques, many focused on the integration of soft and rigid materials for mechanical gradients. While some hybprinters explored droplet printing or multi-material printing for biological gradients, the gradients were mostly limited to a single continuous pattern, limited by the inflexibility in changing materials in single-channel inkjet printing or single-vat DLP-SLA. The capability to generate combinations of biological signals in any desired patterns is yet to be explored.

Our group also joined the effort in investigating hybprinting for biomedical applications. In 2015, our group pioneered integrating DLP-SLA with MME to print vascularized tissue constructs ^44^. Our recent article also reviews various hybprinting techniques for musculoskeletal tissue engineering applications ^45^. In this study, we present our second-generation hybprinter, Hybprinter-SAM, by integrating three distinct printing mechanisms: SE, acoustic droplet ejection (ADE), and MME. While SE and MME are commonly integrated modules for hybprinting soft and rigid biomaterials ^14, 15, 20^, ADE is a specific type of droplet printing mechanism, which focuses the energy of an acoustic wave on a liquid-air interface for droplet ejection ^46, 47^. Previously we have demonstrated that ADE can be used to deposit proteins and growth factors onto a rigid scaffold, maintaining the localization of protein patterns to trigger targeted cellular response ^48^. To our knowledge, this is the first time ADE is integrated into a hybprinting system, thereby expanding the scope of other existing hybprinters. Compared to more commonly used droplet printing technologies, such as inkjet printing, ADE possesses several distinct features: 1) ADE generally provides better resolution with higher reproducibility; 2) being nozzleless, ADE largely reduces shear stress during printing, leading to a gentle printing process with less damage to cells or biomolecules; 3) the design of ADE printer heads allows for reduced cost and easier multi-channel integration with zero carryover, which in turn lead to reduced cross contamination, flexibility and wider application potential, particularly in the field of biomedical engineering ^49–51^. The specific ADE module used in this study is compatible with the standard well-plate format of up to 384 wells and a resolution down to 2.5 nL ^51^, allowing for high throughput capacity of bioreagent depositions. With such capability, we can pattern virtually any custom designed biological signals in a hybprinted scaffold. And combined with MME and SE for rigid and soft material integration, we can fabricate a highly biomimetic tissue engineering construct comprising of biological, mechanical and topographical gradients (**Figure 1b**).

To demonstrate the capability of Hybprinter-SAM, a bone-tendon regeneration application was explored. Bone-tendon injuries account for 30% of all musculoskeletal clinical cases with 4 million new incidences worldwide each year ^52^. Regenerating the bone-tendon interface is arguably the most important yet challenging problem in treating bone-tendon injuries. This is mainly because the bone-tendon interface is a highly sophisticated, functionally graded structure with four distinct zones: bone, mineralized fibrocartilage, fibrocartilage, and tendon, where the properties change drastically from bone to tendon across its length of less than 1 mm ^53–57^. This makes the bone-tendon interface extremely difficult to recapitulate yet makes Hybprinter-SAM a strong contender. Within Hybprinter-SAM, the MME module can print polycaprolactone (PCL) for structural support, SE module can print gelatin/fibrin (gelbrin) hydrogel for stem cell encapsulation and tissue development, and most importantly, ADE can deposit growth factors and crosslinkers to pattern and localize biological signals to guide stem cell development into the bone-tendon interface. Hybprinter-SAM can theoretically print a 3D construct with bone biofactors patterned on the bottom layer, fibrocartilage biofactors on the middle layer, and tendon biofactors on the top layer; however, for simplicity and delivering a proof of concept of stimulating stem cells locally, this study focuses on effecting fibrocartilage markers with FGF-2, a studied growth factor commonly used in chondrogenesis ^58–60^. To carry out the study, the printing process and selected biomaterials were first characterized, including printability, degradation, mechanical property, and biocompatibility. Second, an *in situ* crosslinking strategy via ADE deposition was developed. The crosslinker was printed in a custom pattern to achieve graded crosslinking density, thereby controlling the localization of the biofactors. Lastly, human mesenchymal stem cell (hMSC) differentiation in the hybprinted bone-tendon mimicking construct was studied. Results showed that FGF-2 patterned by ADE module had positive effects in upregulation of fibrocartilage differentiation markers including SRY-box transcription factor 9 (SOX9) and Scleraxis (SCX) ^61–63^, validating Hybprinter-SAM’s ability to pattern and localize biological factors for functional gradients. Additionally, an automated bioreactor was developed to apply mechanical stimuli to the hybprinted scaffold in addition to biological signaling to better mimic in vivo conditions. With such expanded capabilities, Hybprinter-SAM holds promise in a variety of applications where engineering highly sophisticated tissues is needed, and it can serve as a platform technology for further scientific studies in the field of tissue engineering.

## RESULTS

### Hybprinter-SAM system

A novel fabrication system, Hybprinter-SAM (SE+ADE+MME), was developed to engineer multi-material tissue constructs consisting of bioplastics, hydrogels, and liquids by combining three printing mechanisms: syringe extrusion (SE), acoustic droplet ejection (ADE) and molten material extrusion (MME) (**Figure 2**). In terms of hardware integration, the SE module resides on the MME printer head via a custom designed 3D printed adaptor (**Figure 2a**). Therefore, the SE print pathway is easily governed by the same control mechanism of the MME module. The ADE target plate, a built-in feature of the ADE module, was used as the build plate for the whole hybprinting process for ease of integration (**Figure 2a, Figure S1 and Video SV1**). In-house software was developed in C++ to coordinate all three modules, resulting in a fully automated machine that enables hybprinting and seamless integration of three distinct materials under one platform (**Figure 2b, Supplemental Figure S2**). The platform prints in a serial, layer-by-layer fashion. By optimizing and calibrating software and material properties, a layer precision of 100 μm, strut resolution of 150-800 μm, and droplet resolution of 2.5 nL (∼200 μm footprint) were achieved.

**Figure 2.**
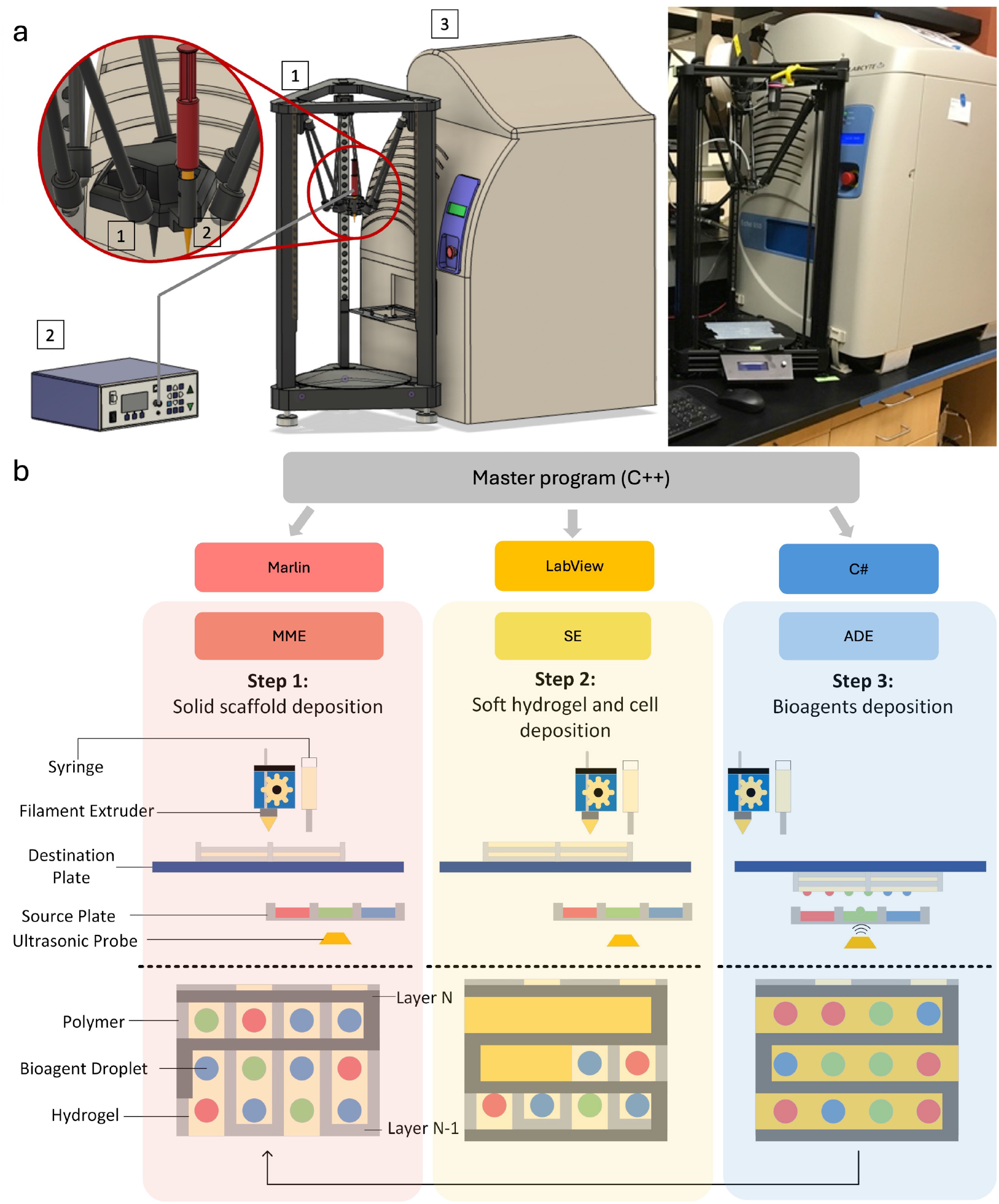
(a) Schematic and assembly of Hybprinter-SAM. (b) Rationale and flowchart for hybrid printing comprising of molten material extrusion (MME), syringe extrusion (SE) and acoustic droplet ejection (ADE). All three modules are coordinated by a custom developed software program, and they follow a layer-by-layer fabrication protocol in the order of MME→SE→ADE, resulting in an engineered construct comprising of rigid polymeric scaffold, soft hydrogel scaffold, and aqueous-phase bioreagents.

The versatility of Hybprinter-SAM was demonstrated via a series of hybprinted scaffolds (**Figure 3, Supplemental Figure S3 and Video SV2-6**). First, the capability of integrating rigid and soft materials with MME and SE modules was demonstrated (**Figure 3a-d, Supplemental Figure S3a-e and Video SV2-4**) by printing different designs of PCL interlacing with hydrogel, resulting in a robust soft-rigid connection. The mechanical integration of the two dissimilar materials was evidenced by the bridge remaining intact while withstanding compression, tension, bending, and torsion (**Figure 3a-d, Video SV2-4**). This integration was further corroborated through destructive testing involving the pulling of a hybprinted structure composed of PCL infilled by hydrogel materials (**Supplemental Figure S3c-d**). The seamless integration of soft and rigid materials in the hybrid scaffold enables it to be rolled up, facilitating improved surgical manipulation, particularly in minimally invasive procedures such as arthroscopic surgery (**Supplemental Video SV5**). Interestingly, the limiting factor is not the soft-rigid interface but the material properties of the hydrogel itself as evidenced by the ruptures within it (**Supplemental Figure S3d**). Secondly, the combination of MME and ADE modules demonstrated Hybprinter-SAM’s ability to precisely deposit biological signals, as represented by food dyes, onto MME printed struts (**Figure 3e-h, Supplemental Figure S3f-g**). Similarly, ADE printed droplets could also be precisely placed onto SE printed hydrogel scaffolds in various patterns, including graded horizontal or vertical patterns, with multiple colors also representing different biochemical cues (**Figure 3i-m, Supplemental Figure S3h-i**). Third, combining all three modules allows for a wide range of designs to be produced, showcasing the versatility of Hybprinter-SAM, such as a Stanford logo print with a base PCL scaffolding and interlacing hydrogel, patterned with graded red dye transitioning from dark (bottom) to light (top) and a green tree layer atop the S fading in the reverse direction (**Figure 3n and Supplemental Video SV1**). While these hybprinted constructs display only a few colors, the ADE module allows for parallel printing of up to 384 different solutions, which facilitates high throughput testing of many combinations of biological/chemical signals. To better demonstrate this capacity visually, a lattice structure with an increased number of colored solutions was printed (**Figure 3o**). Furthermore, to illustrate physiologically relevant designs, a bone-soft tissue mimicking construct was hybprinted with a cylindrical bone mimicking PCL scaffold, and a half-cylindrical hydrogel mimicking soft tissues surrounding the bone (**Figure 3p, Supplemental Figure S3k-l**). Different biological cues can be patterned onto bone or soft tissue scaffolds respectively to induce guided regeneration of bone and soft tissues. The overlapping purple between deposited blue and red dyes also represents a gradient of biochemical cues that can be achieved via hybprinting (**Figure 3p**). A smooth functional gradient was further highlighted with a hybrid bridge structure bookended by PCL lattice squares, where a high concentration of transglutaminase (10 wt/vol%) and thrombin as crosslinkers were deposited by the ADE module onto the hydrogel region. Blue food dye was added to the transglutaminase and thrombin mixture to aid visualization of the graded pattern (**Figure 3q**). As Hybprinter-SAM increasingly deposited more crosslinking factors along one axis onto each layer of gelbrin, the sample predictably showed less stretching (23.6%) in the darker sections (hydrogel struts are closer to each other) than in the lighter sections (67.8%, hydrogel struts have greater gaps) (**Figure 3r and Supplemental Video SV6)**. This demonstrates the capability of Hybprinter-SAM to pattern not only biological factors, but also chemical crosslinkers. Altogether, Hybprinter-SAM’s ability to combine soft and rigid biomaterials, as well as pattern biochemical signals, demonstrate potential for various tissue engineering applications where functionally graded scaffolds are needed.

**Figure 3.**
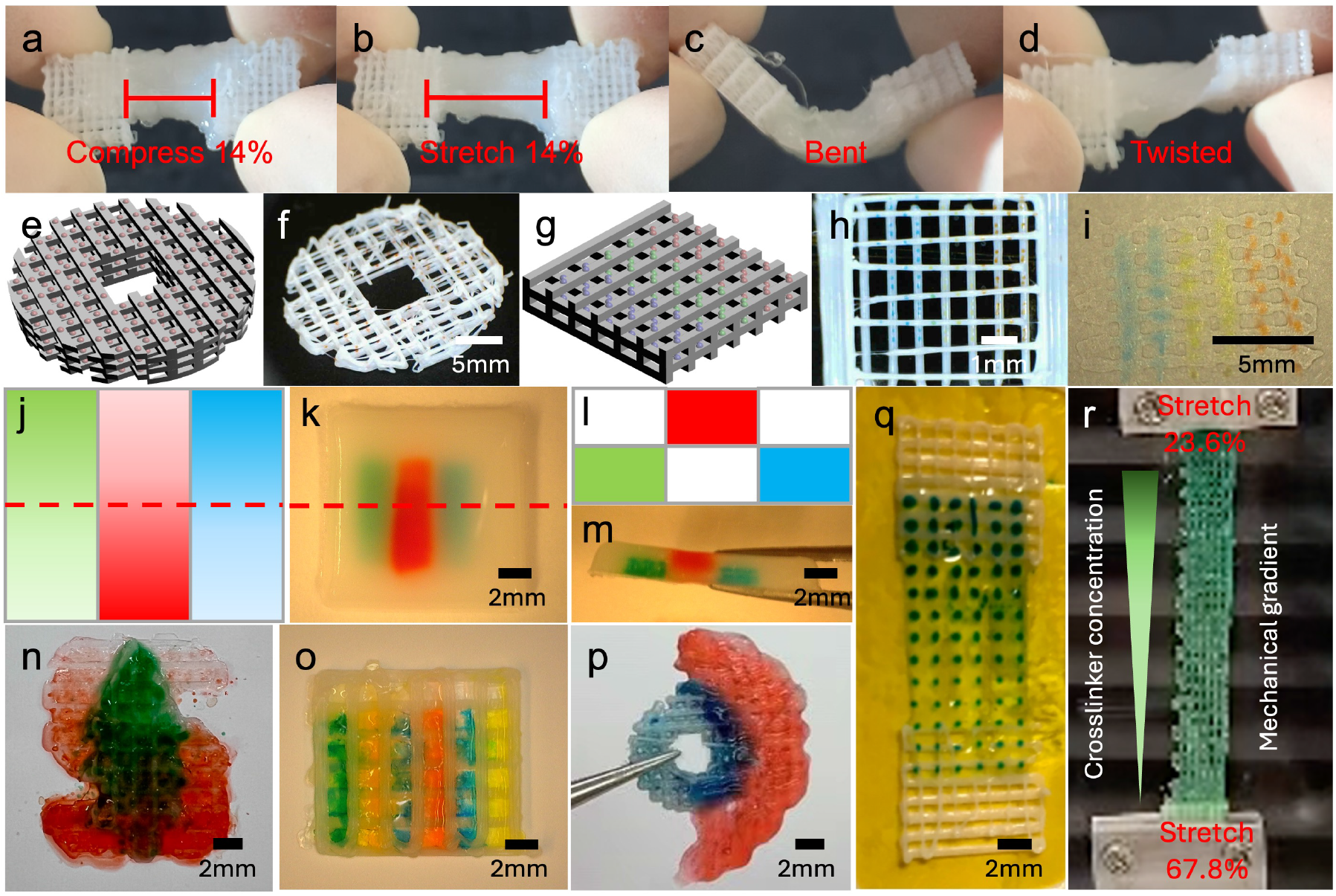
(a-d) Hybrid soft-rigid tissue constructs of hydrogel and PCL that can withstand mechanical manipulation. (e-h) ADE deposition of droplets onto MME printed PCL struts. (e,g) Conceptual design. (f, h) Printed with Hybprinter-SAM. (j-m) Hydrogels with biological signal patterns in both horizontal and vertical directions. (j, l) Conceptual design. (k, m) Printed with Hybprinter-SAM. (j, k) Top view. (l, m) Cross-sectional view. (n-p) Soft-rigid hybrid constructs with multiple biological factors (demonstrated by dyes) with different patterns, enabled by Hybprinter-SAM. (q-r) Biological/chemical gradient across the scaffold, demonstrating a mechanical gradient (r).

### Optimization and characterization for bioink materials and bioprinted scaffolds

To cover a broad range of potential applications, as well as to optimize for the specific proof-of-concept application, polycaprolactone (PCL) bioplastic, gelbrin hydrogel and aqueous solutions of growth factors or crosslinkers were selected in this study as the printing materials for MME, SE and ADE modules, respectively. Both PCL and gelbrin are established biomaterials for various bioprinting applications ^64–69^. The printing process of PCL was systematically optimized by tuning parameters such as printing temperature, extrusion rate, and printer head movement speed in an open-source slicing software (UltiMaker Cura). Optimized parameters are listed in **Supplemental Figure S4**.

Gelbrin hydrogel was utilized in a modified formulation originally developed by D. Kolesky et al. ^69^ due to its excellent biocompatibility, attributed to its two naturally derived components: gelatin and fibrin. In brief, the bioink preparation employed the same gelbrin composition (7.5% gelatin, 1% fibrinogen), but the crosslinking solution was modified to consist of 50 units/mL thrombin, 12.5 mM CaCl_2_, and 1.5 wt/vol% transglutaminase (TGase). The temperature-dependent physical gelation of gelatin ensured the printability of the gelbrin bioink. Upon printing, samples were submerged in the crosslinking solution, where fibrinogen rapidly crosslinked into fibrin, subsequently followed by Ca^2+^ dependent enzymatic crosslinking of gelatin and fibrin by TGase (**Supplemental Figure S5**). The properties of the modified gelbrin hydrogel, both as a bioink prior to crosslinking and as a construct post-crosslinking, were optimized and characterized through a series of studies. Before crosslinking, the rheological property of the gelbrin bioink was characterized to assess its printability. An oscillation test with temperature sweep was conducted to evaluate the temperature-dependent rheological property of gelbrin (**Figure 4a**), primarily influenced by the gelatin component which liquifies at 37 °C. Based on the results of the temperature sweep, 20 °C was selected as the printing temperature, ensuring stable physical gelation necessary for printability. Further validation of printability was conducted using a rotational shear test as shown in **Figure 4b**. The results indicated a decrease in viscosity with increasing shear rate, confirming the shear-thinning property of the gelbrin bioink. This shear-thinning behavior is crucial for bioinks used in SE printing, as the bioink needs to be sufficiently non-viscous enough for printing, while maintaining adequate viscosity to uphold its own structural integrity post-printing. Collectively, these results demonstrate the printability of gelbrin bioink at 20 °C, which was maintained as the environmental temperature throughout the process.

**Figure 4.**
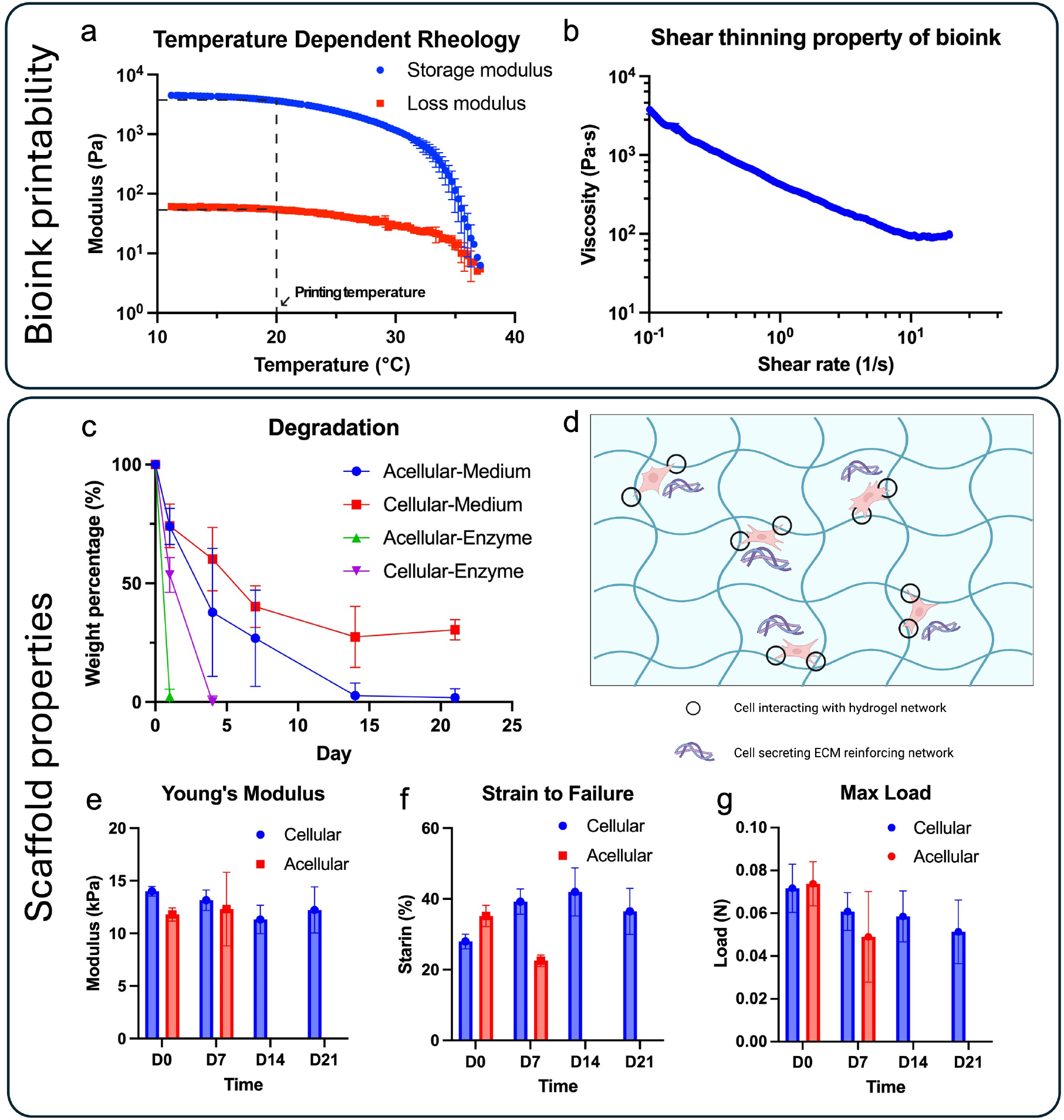
(a-b) Characterization of the bioink printability. (a) Temperature-dependent rheological property of the bioink. (b) Shear-thinning property of the bioink. (c-g) Characterization of the scaffold after printing and crosslinking. (c) Degradation profile of the hydrogel in culture medium or 0.1% collagenase solution. (d) Possible mechanisms causing slow degradation in cellular samples. (e-g) Mechanical properties of the printed samples. (e) Young’s modulus of printed samples in a tensile test. (f) Maximum strain to failure of printed sample. (g) Maximum load of the printed sample.

After printing and crosslinking, the degradation and mechanical properties of gelbrin hydrogel were quantified. The degradation profile included an evaluation of both the intrinsic degradation of acellular gelbrin and the degradation of hMSC-laden gelbrin. The latter assessment more accurately reflected the conditions of subsequent applications, where hMSCs were loaded and studied for bone-soft tissue regeneration. Degradation was profiled under both normal medium and accelerated enzymatic (0.1% collagenase) conditions. Accordingly, the acellular gelbrin samples degraded at an accelerated rate compared to the cellular samples under both regular medium and enzymatic conditions as shown in **Figure 4c**. As the root of this disparity, we hypothesize that the hMSCs either interacted with the hydrogel matrix by grasping onto the gelbrin material, generated their own extracellular matrix, or secreted crosslinking enzymes to reinforce the hydrogel as they colonized it ^70–73^ (**Figure 4d**), thereby slowing down degradation. Notably, shrinkage of the gel was observed during the incubation (**Supplemental Figure S6**), providing further support for the first hypothesis that the hMSC interactions with gel may have exerted a pulling force, contributing to the observed shrinkage.

The results align with the findings on the mechanical properties of both acellular and cellular gelbrin (**Figure 4e-g**). Note that mechanical properties were measured in water bath (**Supplemental Figure S7a and Video SV7**), as the gel would deform and affect the measurement when measured in air (**Supplemental Figure S7b-c**). The mechanical strength of acellular samples declined in the first week, with complete degradation occurring by Week 2, resulting in no available samples for mechanical measurement (**Figure 4c and Supplemental Figure S6**). In contrast, cellular gelbrin preserved its mechanical strength over the course of 3 wks. The slight decrease in Young’s Modulus of the cellular gelbrin over the first 2 wks may be attributed to the degradation of the gelbrin, while the increase observed past Week 2 could be a result of cellular influences over time. Moreover, the strain to failure increased largely for the cellular sample, potentially due to hMSCs reinforcing the hydrogel network, whereas the strain data for acellular samples exhibited a marked decrease (**Figure 4f**). Interestingly, the maximum load continued to decrease rather than following the same trend as strain to failure, likely due to gel shrinkage induced by the presence of cells, leading to a reduced cross-sectional area and, consequently, lower load capacity (**Figure 4g**).

### Hybprinting of cell-laden bone-tendon constructs

As a proof-of-concept application, an hMSC-laden bone-tendon construct was designed to illustrate the potential of Hybprinter-SAM for tissue engineering. Bone-tendon was selected as the target application due to its complex region with graded compositional, mechanical, biological properties ^53–57, 74–77^. For the purposes of this application, the cytocompatibility of the materials was validated. Initially, hMSCs were encapsulated in the hydrogel and printed via the SE module. The cell viability was evaluated through a live and dead assay, which indicated > 95% viability 3 weeks post-printing (**Figure 5a-b**). Although there was a slight decrease in viability on D4, which dropped to around 80%, this decline is associated with gel degradation at the edges as most of the dead cells were observed along the periphery of the hydrogel. We hypothesize that this is due to accelerated degradation of gel along the edges or more exposure to the crosslinking solution. Encouragingly, the cells remained healthy in the majority of the gelbrin, exhibiting elongation and connection to neighboring cells on D4 (**Figure 5a**). These findings demonstrate the biocompatibility of the hydrogel material.

**Figure 5.**
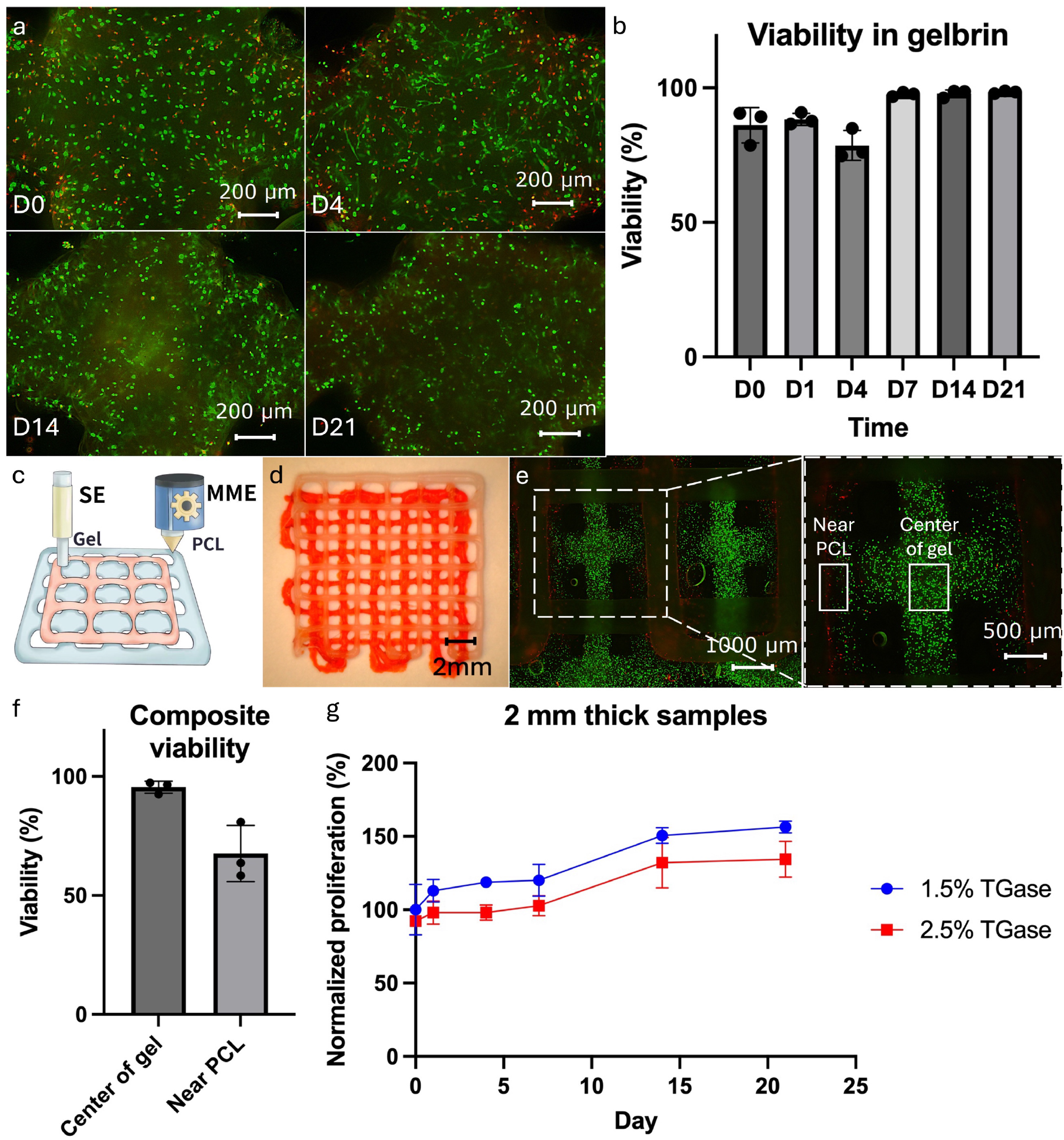
(a-b) Cell viability study for the hydrogel material. (c-f) Cell viability study for the composite PCL and hydrogel printed sample. (c) Schematic design and (d) printed sample of the complementary pattern with PCL and hydrogel. (e) Fluorescent picture and (f) quantified viability of the composite sample. (g) MTS study for proliferation of the printed cell-laden hydrogel sample.

The biocompatibility of the materials undergoing the hybprinting process was further validated, as the MME module operates under high temperatures, which could adversely affect cell viability. To assess this, composite samples composed of PCL and hMSC-laden gelbrin were printed via the MME and SE modules. The pair of rigid and soft materials were printed in a complementary pattern as shown in **Figure 5c-d**, where the two paths intersect during printing. The cross-sections were of particular concern as the high-temperature PCL strut would make direct contact with the cell-laden hydrogel. Viability analysis was performed post-print (**Figure 5e-f**), which demonstrated that in the bulk of the hydrogel, cells remained highly viable (>95% viability), with only cells within 200 μm of the PCL strut being affected (68% viability). This can be attributed to the rapid heat dissipation of the hydrogel material. This phenomenon was further validated in a COMSOL simulation (**Supplemental Figure S8**), which shows the hydrogel region within 300 μm of the MME printed strut reaching high temperatures before quickly dropping and remaining below body temperature (37 °C). The simulation is consistent with the experimental data and collectively affirms that the entire printing process minimally affects cell viability.

Cell survival in long-term cultures of the printed constructs was also evaluated. Constructs of two thicknesses (1 and 2 mm) were printed and assessed over 3 wks using MTS assays, which measure the metabolic activity of the hMSCs. An increase in the metabolic activity was detected, indicating cell proliferation (**Figure 5g and Supplemental Figure S9**). Notably, thinner samples and lower TGase concentrations resulted in faster proliferation rates, suggesting improved nutrient exchange in these groups. Overall, the biocompatibility of the Hybprinter-SAM printing process and the biomaterials used was validated, laying the foundation for the remainder of our bone-tendon tissue engineering study.

### Development of fibrocartilage for bone-tendon regeneration

Fibrocartilage differentiation was investigated as the next step in the bone-tendon tissue engineering application. To induce differentiation, the ADE module was used to pattern FGF-2 on the hybprinted construct to guide differentiation into fibrochondrocytes. There are several growth factors reported to have potential in promoting fibrochondrocyte differentiation, including TGF-β3, FGF-2, PDGF-BB, etc. ^53,78^ Among them, FGF-2 is known for promoting bone-tendon healing by multiple means, including enhanced cell proliferation and differentiation ^58–60^.

Before assessing the impact of FGF-2 specifically, fluorescein isothiocyanate (FITC)-conjugated bovine serum albumin (BSA) was selected as a model protein to quantify the drug loading and release kinetics of gelbrin, as BSA is commonly used in similar studies ^79^. Two main aspects of drug loading and delivery were quantified: desorption from the hydrogel and diffusion within the hydrogel, akin to ink spreading on paper. BSA was patterned via ADE onto the hydrogel scaffolds (**Supplemental Figure S10d**) with individual droplets (10, 50, and 100 nL per spot) creating distinct lines. These varying volumes corresponded to total amounts of 16.2, 81, and 162 μg of BSA in each scaffold, respectively. The scaffolds were then crosslinked for 1 hr and submerged in PBS. At each time point, PBS was collected and analyzed for fluorescence desorption, while the scaffolds were imaged for fluorescence retention. This protocol resulted in a burst release that resulted in virtually no long-term protein retention: about 80% of the loaded protein diffused out during crosslinking (**Figure 6e**). To achieve a controlled release and localization of biofactors, the protocol was modified to dissolve BSA into thrombin prior to loading into the ADE, enabling *in situ* crosslinking during the print process. The rapid crosslinking between fibrinogen and thrombin at each printed layer created a localized high-density crosslinking network which effectively locked in the BSA, resulting in the expected slower release profiles and longer retention (**Figure 6a**). Results show that the printed pattern remained visually distinct after a 3 wk culture (**Figure 6b**) indicating minimal diffusion within the hydrogel. Additionally, both the low (10 nL) and high (50 nL) initial BSA loadings demonstrated a sustained release throughout the course of the study (**Figure 6c and Supplementary Figure S10b**), and correspondingly, the integrated density of fluorescence on the patterned halves stabilized after a few days (**Figure 6d**). The high initial loading resulted in a reduced desorption rate, likely due to a minimum quantity near the hydrogel surface desorbing in both groups but representing a smaller percentage in the higher loading group. Fluorescence quantification from both the unprinted and printed halves support this, as the slopes from D0 to D1 are similar for both loading groups (**Figure 6d**). Additionally, images of the scaffolds show that the BSA that was desorbed from the hydrogel was not re-adsorbed, as supported by the constant fluorescence intensity at the unprinted halves. Similar results can be achieved by printing the desired protein first, followed by thrombin (**Figure 6a**), the flexibility of which is a byproduct of the ADE module.

**Figure 6.**
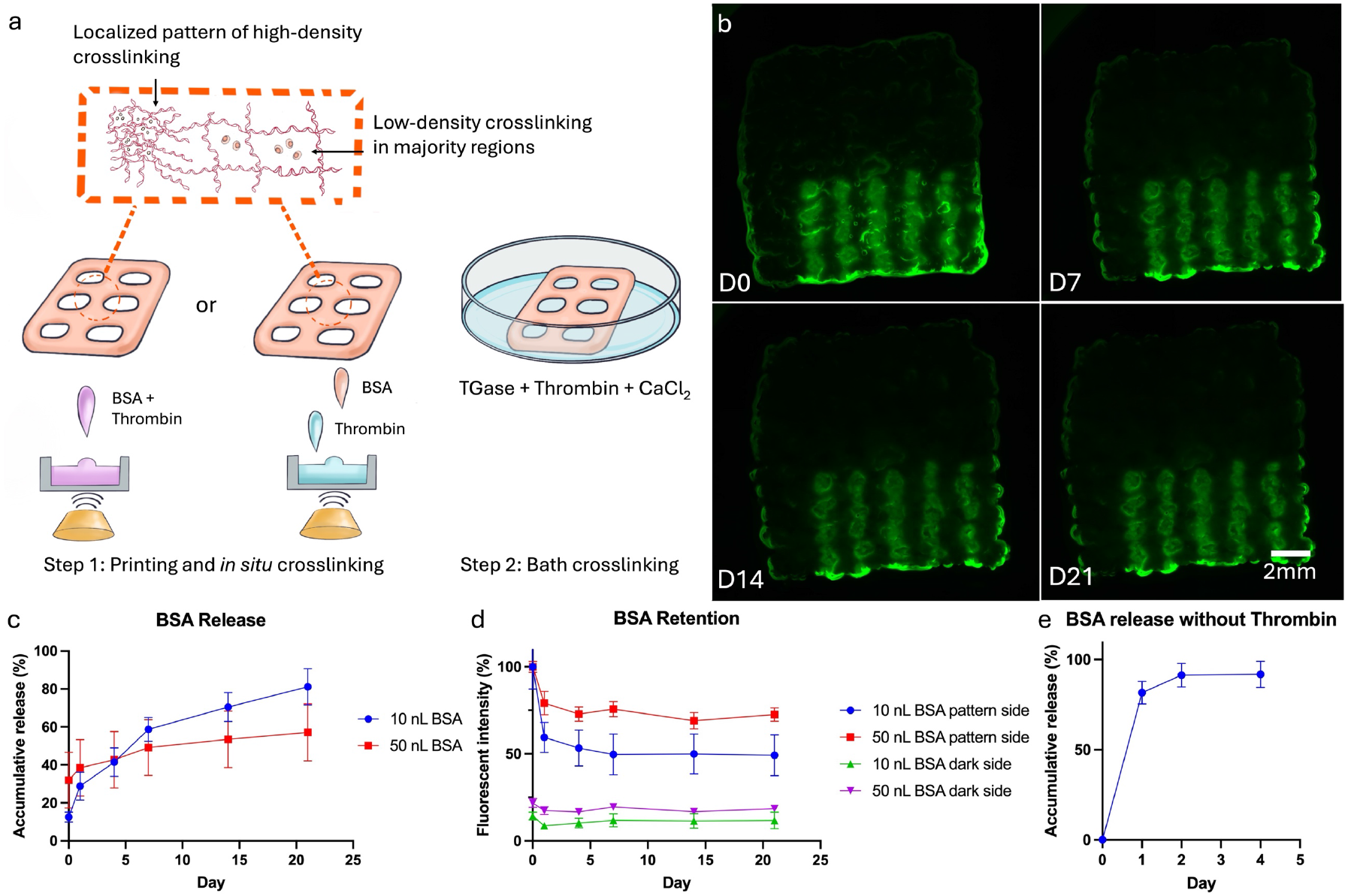
(a) In situ crosslinking strategy for protein immobilization in the gelbrin dual-crosslinking network. (b) Fluorescent images showing controlled release and pattern retention of FITC-BSA printed in droplets of 50 nL. (c-d) Quantified BSA release and retention of in situ crosslinked samples. (e) BSA release of non-in situ crosslinked sample.

Note that while the release and retention kinetics for both acellular and cellular samples were quantified (**Supplemental Figure S10a**), acellular conditions better reflect the intrinsic retention capability of the material itself, as it rules out the possibility of cellular uptake of proteins of interest. But as previously mentioned, acellular samples have accelerated degradation at 37 °C relative to cellular samples, so to ensure the integrity of the acellular samples, the release studies were performed at room temperature. Additionally, even higher BSA loadings (100 nL) were investigated and demonstrated a saturation point where sufficient BSA desorption and re-adsorption occurred, resulting in visible fluorescence on the unprinted halves (**Supplemental Figure S10c**).

Utilizing this controlled localization of biological cues via *in situ* crosslinking, the effects of patterned FGF-2 on hMSCs were studied. A bone-tendon mimicking scaffold was hybprinted with PCL modeling bone and gelbrin modeling tendon, along with FGF-2 patterned uniformly via ADE (**Figure 7a)**. Specifically, FGF-2 was printed at a concentration of 5 μg/mL in the soft hydrogel region in an array format with 15 nL per spot. After incubation in chondrogenic medium for up to 7 days, the hMSC differentiation was evaluated using quantitative polymerase chain reaction (qPCR) to assess gene expressions of *SCX, SOX9, COL1, COL2* and *AGG. SCX* and *SOX9* are early-stage markers for fibrocartilage differentiation; *SCX+* cells tend toward tendinogenic differentiation, while *SOX9+* cells lean toward chondrogenic and osteogenic differentiation ^61–63, 80^. *COL1, COL2* and *AGG* are later stage markers indicating extracellular matrix deposition, with collagen type 1 (*COL1*) present in all zones (bone, fibrocartilage, and tendon), collagen type 2 (*COL2*) mainly in fibrocartilage, and aggrecan (*AGG*) also in fibrocartilage ^53, 57, 74, 81^.

**Figure 7.**
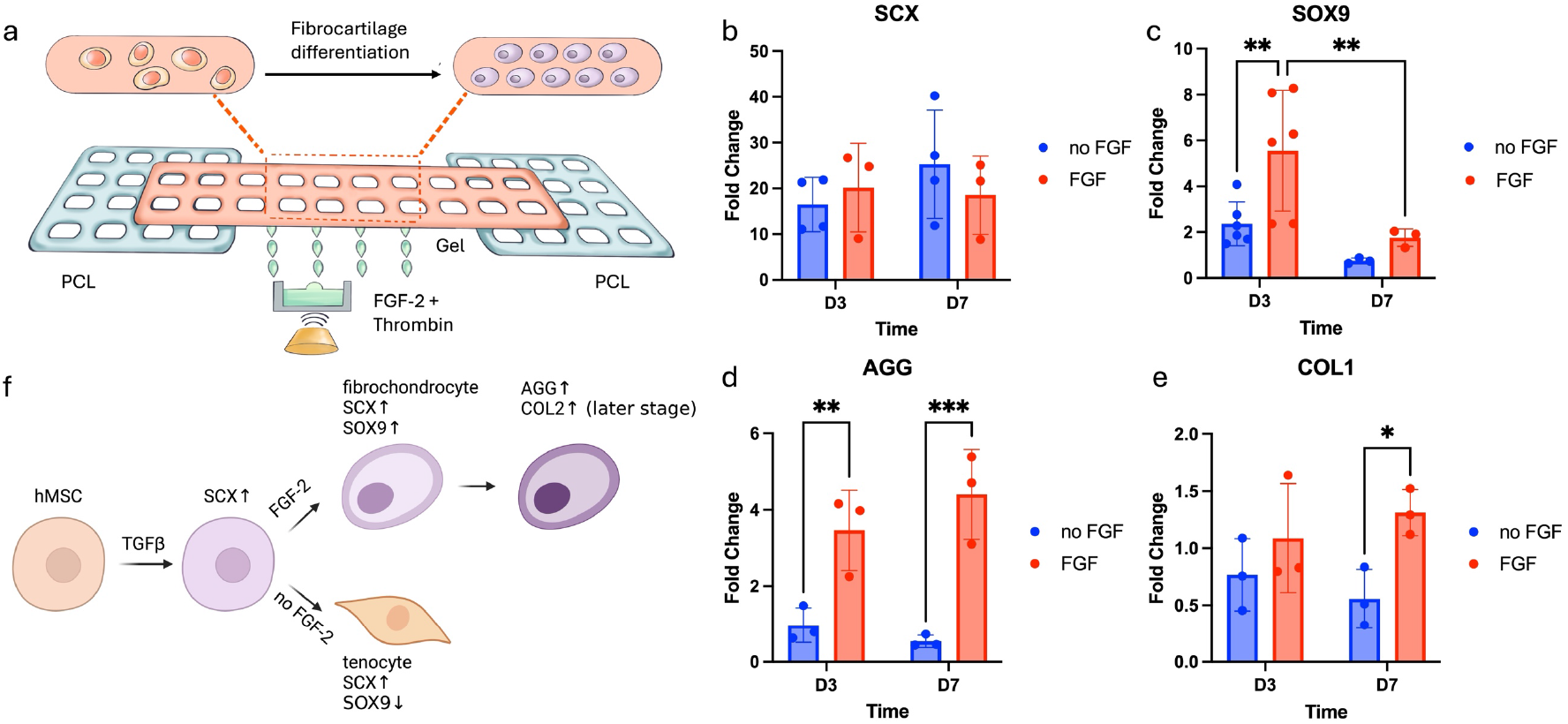
(a) Schematic of the bone-tendon mimicking construct with biological gradient. (b-e) qPCR expression of (b) SCX, (c) SOX9, (d) COL1 and (e) AGG for the study of hMSC differentiation in the bone-tendon mimicking construct. (f) Possible pathways for fibrocartilage differentiation.

Cells encapsulated in hydrogel with and without FGF-2 were analyzed. Scaffolds were cultured for three or seven days, and gene expression was compared to Day 0 levels. Results showed significant upregulation of *SCX* across all groups, likely due to the chondrogenic medium (**Figure 7b**). *SOX9* was upregulated only in the FGF-2 patterned group (**Figure 7c**), indicating that FGF-2 was the primary factor. Upregulation of *AGG* was also observed, suggesting differentiation into fibrocartilage (**Figure7d**).

No expression of *COL2* (data not provided) was observed and *COL1* remained almost constant (fold change ∼1) across all groups (**Figure 7e**), likely because cells were still in early differentiation stages without significant extracellular matrix deposition. These findings align with existing studies ^53, 82–84^ and further validate the role of FGF-2 in promoting fibrocartilage differentiation. TGF-β3 in the chondrogenic medium may have acted as an upstream differentiation factor which leading to upregulation of *SCX* ^85, 86^, while FGF-2 may have promoted upregulation of *SOX9*, further driving fibrocartilage differentiation ^53, 84^ (**Figure 7f**). Without FGF-2, cells may have undergone tendinogenic differentiation.

To further investigate bone-tendon regeneration, mechanical stretch was applied to the hybprinted bone-tendon mimicking scaffold, as studies have shown that mechanical stimuli play important roles on the development of musculoskeletal tissues, particularly at bone-tendon interfaces ^87–89^. A custom designed mechanical bioreactor was manufactured to apply tension or compression to up to 6 samples simultaneously in a 6-well plate(**Supplemental Figure S11a-d and Video SV8**). The bioreactor can be sterilized, fit within a standard CO^2^ incubator, and can provide diverse loading conditions that mimic physiological mechanical stimuli (static stretch, slow stretch, cyclic tension, cyclic tension and compression) (**Supplemental Table S12**) ^87–89^. For simplicity, only slow stretch tension was applied to the hybprinted samples with and without patterned FGF-2 and were evaluated for hMSC differentiation via qPCR. Results did not show significant differences in *SOX9* or *AGG* expression between samples with and without tension (**Supplemental Figure S11e-f**); however, a slight increase in *SOX9* expression was observed in samples with FGF-2 under tension, and a slight increase in *AGG* expression in samples without FGF-2 under tension. While further research is needed to understand the combined effects of mechanical stimuli and biological gradients on fibrocartilage differentiation (**Supplemental Figure S11g**), Hybprinter-SAM offers the potential to systematically study the interplay of mechanical, biological, and material factors in a high-throughput manner. Overall, the results validate the use of patterned biological signals with Hybprinter-SAM to induce hMSC differentiation, demonstrating potential in engineering complex, functional tissues.

## DISCUSSION

Despite rapid advancements in tissue engineering, there remains a significant need for advanced technologies capable of manufacturing tissues and organs that closely mimic their native counterparts. We propose hybprinting as a prospective solution to this challenge. Our research has focused on combining multiple printing mechanisms to overcome limitations inherent in single-method printing techniques ^44, 45^. Notably, numerous research groups have begun exploring hybprinting approaches, underscoring its importance and substantial potential in the field ^14–21, 31–40^. To date, most studies have combined up to two different printing mechanisms. A common strategy involves using separate modules to address soft and rigid materials, or employing a secondary printing module to assist the primary one ^14–21, 31–40^. While some emerging technologies are investigating the integration of more than two printing mechanisms ^41–43^, this area remains relatively unexplored due to the complexities associated with system integration and subsequent application studies.

In this study, we combined three distinct printing mechanisms – SE, ADE, and MME – to develop Hybprinter-SAM. This integration significantly expands our capability to fabricate complex scaffolds, and to the best of our knowledge, this is the first instance where ADE has been introduced into a hybprinting system. The inclusion of ADE provides a critical feature: versatile and precise patterning of biological and chemical cues onto engineered scaffolds. This was specifically achieved by the multi-well plate ADE system, compatible with up to 384-well plates, which offers an almost unlimited capacity for patterning combinatorial signals – including proteins, small molecules, cells, and crosslinkers – without the need to change plates or nozzles. By integrating SE and ME with ADE, we have demonstrated the ability to produce improved native-mimicking scaffolds that exhibit topographical, mechanical, and biological gradients in a seamless, integrated process (**Figure 1**).

An optimal bioink or biomaterial is fundamental to the success of bioprinting applications; however, the ideal bioink may vary depending on specific applications ^90, 91^. Hybprinter-SAM has demonstrated versatility in processing a wide range of biomaterials, including rigid thermoplastics, soft hydrogels, and biochemical reagents; it accommodates materials with mechanical properties spanning over 7 orders of magnitude. In this study, the biomaterials were optimized for a bone-tendon modeling application as a proof of concept. Specifically, our group has had extensive experience with PCL-based materials for bone tissue engineering ^64–67^, as well as gelatin or GelMA-based hydrogels for cell encapsulation and delivery of bioactive signals ^92–94^. And accordingly, a gelatin/fibrin hybrid material was chosen as the hydrogel carrier for cells and bioactive signals. Originally developed by D. Kolesky et al. for its excellent biocompatibility ^69^, the fabrication protocol was modified to improve handling and printability by tuning the crosslinking bath formulation and duration and printing parameters such as temperature, nozzle sizes and printing pressure.

The bioink rheology studies demonstrated that controlling the printing temperature facilitated desired shear-thinning properties for enhanced printability (**Figure 4a-b**). Combined with mechanical and degradation characterizations, as well as viability and proliferation studies, the hydrogel system was optimized to balance printability, structural integrity and biocompatibility (**Figure 4c-g**). Achieving such a balance is a persistent challenge in bioprinting as improving mechanical strength often opposes cytocompatibility ^95–97^. Various studies have explored strategies to attain this balance, including leveraging the stress relaxation property of hydrogel post-crosslinking ^98, 99^, designing micro-porous hydrogels ^100, 101^, or utilizing multiple crosslinking mechanisms ^102–104^. The gelbrin system employs a tri-crosslinked network among temperature-dependent physical gelation of gelatin, chemical crosslinking of fibrinogen by thrombin, and gelbrin by TGase ^69^ (**Supplemental Figure S5**). This hierarchical crosslinking approach enables printability and desired mechanical properties at different stages through varying degrees of crosslinking. Furthermore, as all components of the gelbrin system are naturally derived polymers, it ensures biocompatibility, facilitating cell-hydrogel interactions and allowing cells to gradually digest the hydrogel, while producing their own ECM. The cell-hydrogel interaction was observed through contraction of the cell-laden hydrogel, resulting in a slower degradation. However, the underlying mechanisms of these cell-hydrogel interactions and the potential role of cell-secreted ECM in reinforcing the hydrogel warrant further investigation in future studies.

Controlled localization of biological signals within the hybprinted scaffold is essential to the success of the proposed research. Without it, even though Hybprinter-SAM can print biological patterns on the scaffold, the molecules may diffuse out prior to their signals having any desired effect. To address this, an *in situ* crosslinking strategy was developed, involving simultaneous printing of crosslinkers along with biological cues. This strategy for immobilizing proteins is largely enabled by Hybprinter-SAM, as it allows for parallel printing of both biological signals and crosslinkers. With this capability, controlled protein localization was achieved, quantified by both the release of protein into the culture medium and the retention and diffusion within the hydrogel. Moreover, this demonstrates a hydrogel system that exhibits both cytocompatibility and effective protein retention. Oftentimes, these two factors are in opposition: to immobilize proteins, a hydrogel network must be sufficiently dense, which limits cell movement and proliferation ^95, 97, 105^. Alternative solutions to address this trade-off include designing different binding strategies for proteins while maintaining a relatively loose network ^106–108^. In these studies, fibrin provides heparin binding sites for FGF-2, which is also applicable to the gelbrin system. While the crosslinking bath enables multiple crosslinking strategies, including the formation of fibrin from fibrinogen through the action of thrombin, this was insufficient for maintaining long-term retention of patterned proteins in a more general application, such as BSA (**Figure 6e**). Note that BSA also interacts with gelbrin through both the gelatin and fibrin components ^109–111^; however, the network density used in this study was not enough to maintain controlled localization (**Figure 6e**). Therefore, a more rapid, albeit partial, *in situ* crosslinking was added to the print process. The ADE module enabled *in situ* crosslinking to occur only on the precise protein locations, preserving the remaining hydrogel from additional crosslinking to reduce potential adverse effects on biocompatibility (**Figure 6a-d**). And notably this added step was sufficient to immobilize the protein enough to last the duration of the bath for further crosslinking and result in a hybprinted scaffold with a long-term release profile of 3 weeks.

Hybprinter-SAM is well-suited for applications requiring multiple types of localized biological signals, such as high-throughput 3D drug screening or complex tissue modeling ^10, 112, 113^. In these applications, if tens or hundreds of biomolecules need to be patterned and localized, conventional methods – such as embedding biomolecules in bioinks for SE printing ^114, 115^ or using single- or multi-channel droplet printing module ^116, 117^ – offer limited throughput. In contrast, Hybprinter-SAM can pattern a large array of biological cues along with corresponding crosslinkers. Additionally, the degree of crosslinking can be tailored for each molecule, facilitating distinct release profiles for temporally sensitive studies, including multi-stage differentiation.

This capability to maintain biological factors within the hybprinted scaffold enabled the effective patterning of FGF-2 to study fibrocartilage differentiation for a bone-tendon repair proof-of-concept.

Fibrocartilage development was selected due to its relevance in addressing challenges within bone-tendon regeneration. The bone-tendon interface exhibits mechanical and compositional gradients with drastically changing properties within 1 mm ^53–57^ posing two main challenges in regeneration: (1) the underlying mechanisms of fibrocartilage development remain largely unexplored, and (2) engineering a scaffold that mimics the native bone-tendon interface is technically complex. This study focused on addressing the first challenge by utilizing Hybprinter-SAM to model fibrocartilage differentiation *in vitro*. We validated that the appropriate application of growth factors, including TGF-β3 and FGF-2 play important roles in fibrocartilage development (**Figure 7**). Additionally, an automated mechanical bioreactor was developed to apply mechanical stimuli to the hybprinted samples, to better mimic *in vivo* conditions. This study capitalizes on the functionalities of Hybprinter-SAM to engineer a hybrid scaffold with different growth factor combinations (protein localization) as well as rigid-soft materials integration to mimic bone-tendon (mechanical gradient) and withstand mechanical stimuli (external loading) (**Supplemental Figure S11g**). To our knowledge, a systematic study addressing these three factors simultaneously has not been previously reported and could provide valuable insights into the mechanisms of fibrocartilage development. With the capability of Hybprinter-SAM, as already demonstrated by graded color pattern (**Supplementary Figure S3m**), further study can expand into deposition of graded pattern of growth factors for more in-depth understanding of their effects on fibrocartilage differentiation and bone-tendon transition. Future studies may also include development of scaffolds compatible with arthroscopic surgery to be implanted for *in vivo* treatment for bone-tendon repair.

## CONCLUSION

Despite significant advancements in 3D bioprinting, replicating the complexity of native tissues remains challenging due to the inherent limitations of individual printing mechanisms. To address this, we introduced Hybprinter-SAM, which combines SE, ADE and MME, to expand the capability of fabricating complex scaffolds with mechanical and biological gradients. To demonstrate its capabilities, hybrid constructs were printed with physical integration of soft and rigid materials, along with combinations of biochemical signals. Characterizations of the hybprinted scaffolds were conducted to assess and optimize printability, mechanical property, degradation, and biocompatibility. And notably, the inclusion of the ADE module enabled extensive patterning capabilities for biological cues. Additionally through this module, *in situ* crosslinking allowed controlled immobilization of printed biological patterns. As a key application, Hybprinter-SAM fabricated a hybrid scaffold laden with hMSCs and FGF-2 to guide differentiation into fibrocartilage with qPCR results confirming successful early-stage differentiation with upregulation of *SCX, SOX9* and *AGG*. These results underscore Hybprinter-SAM’s ability to engineer constructs integrating dissimilar materials to produce mechanical gradients and pattern biochemical cues. Overall, this platform presents a tool for high-throughput screening and systematic investigation of these factors.

## METHODS

### Gelbrin bioink preparation

The gelatin-fibrinogen (gelbrin) bioink was prepared via a modified protocol first introduced by D. Kolesky et al. ^69^. Briefly, a 15% gelatin (type A; 300 bloom from porcine skin; Sigma, MO) stock was prepared by dissolving in PBS (1xPBS without calcium and magnesium) at 70 °C while stirring for 12 hours. The pH of the gelatin solution was adjusted to 7.5 using 1M NaOH. The stock was then sterile filtered (0.22 μm, Merck, NJ) and aliquoted for storage at 4 °C for later use (<3 mo.). Immediately before printing, the gelatin aliquot was incubated at 37 °C for 45 min. Simultaneously, fibrinogen Type 1-S (20 mg/mL, Sigma, MO) was dissolved in PBS and incubated at 37 °C for 45 min. The incubated precursor solutions were then mixed 1:1 for final concentrations of 7.5% gelatin, 10 mg/mL fibrinogen. This bioink was cooled at 4 °C for 20 min and brought back to room temperature for 15 min before printing at an environmental temperature of 20 °C. The bioink would be printable for 2 hrs.

### Hybprint-SAM setup and hybrprinting process

The three modules of Hybprinter-SAM, including (1) MME (ATOM; Taipei, Taiwan), (2) SE (Nordson EFD; Westlake, OH), and (3) ADE (Labcyte Echo; Indianapolis, IN), were calibrated for printing bioplastics, hydrogels and liquids, respectively. Specifically, MME was utilized to extrude PCL filaments (50kDa, Facilan™ PCL 100, 3D4Makers, Netherlands) at 120 °C, printing strut size of 150-550 μm; SE was used to dispense gelbrin hydrogel at a pressure ranging from 10 to 20 psi, with strut size ranging from 500-800 μm; ADE deposited droplets of crosslinkers of growth factors, with a resolution of 2.5 nL. Prior to printing, PCL bioplastic and gelbrin bioink were loaded onto the MME and SE module respectively, while biochemical reagents were loaded into 384 well plates for ADE printing. Related printing parameters can be found in **Supplemental Table S4**. A custom program (C++) was developed to coordinate all three modules to deposit materials in a serial, layer-by-layer fashion (**Figure 2b**). The master program received the hybprinting designs in G-code, generated by an open source slicing software (Cura) from CAD designs, and parsed the G-codes to feed separate instructions to each module to execute the printing process (**Supplemental Figure S2**).

Upon printing, the tissue construct was immediately crosslinked for 1 hr at room temperature by the crosslinking solution, composed of thrombin (bovine, Sigma, MO), CaCl_2_ (Thermal Fisher Scientific, MA) and transglutaminase (TGase Moo Glue, TI formula). Stock thrombin was prepared at 500 units/mL and frozen at −80 °C. Aliquots of the stock were thawed right before use. Stock CaCl_2_ solution in PBS was prepared at 125 mM and stored at room temperature. TGase was stored in powder and dissolved right before use. The final crosslinking solution was composed of 50 units/mL thrombin, 12.5 mM CaCl_2_, and 1.5 wt/vol% TGase in PBS. Crosslinked samples were then washed with PBS and incubated in cell culture medium or PBS for further study.

### Rheology Measurement for the Bioink

Gelbrin hydrogel bioink was prepared and stored in a syringe at 4 °C. Rheometer (ARES-G2, TA Instrument, DE) was used to measure the bioink’s rheological property. Upon measurement, 0.5 mL of the bioink was manually extruded onto the test plate of the equipment and sandwiched between the test plate and a 40 mm cone-shaped plate with 0.1 rad angle. Extra gel was removed with a spatula before to make sure gel only remained in the sandwiched part. Two tests were performed, including an oscillation test with temperature sweep from 10 °C to 37 °C, to measure the storage and loss moduli of the bioink, as well as a rotational test at 20 °C with shear rate increased stepwise from 0.1-20 1/s to measure the viscosity change under different shear rates.

### Degradation profile of gelbrin

Bioink was prepared and printed into 2 cm × 2 cm × 2 mm mesh samples, with strut pitch-to-pitch distance of 2 mm. After printing and crosslinking, cellular or acellular samples were cultured in Dulbecco’s Modified Eagle Medium (DMEM) (Thermal Fisher Scientific, MA) or 10 μg/mL collagenase for up to 3 wks. Samples were collected at different time points and their weights after freeze drying were recorded and normalized to weight at D0 for quantification of the degradation.

### Mechanical Characterization

Samples were hybprinted into a gelbrin lattice bookended by PCL lattices. Tensile tests were performed for the hybprinted samples using a universal mechanical testing platform (Instron 5944, Instron Corporation; Norwood, MA) fitted with a 100 N load cell (Interface Inc.; Scottsdale, AZ). For measurement, samples were submerged in a water bath at 37 °C, gripped by two ends and stretched at 1%/sec until failure. A preload of 0.1 N was applied at the beginning of each measurement. The strain and tensile stress were recorded for each measurement to calculate the Young’s Modulus and maximum load.

### Cell printing and biocompatibility study

hMSCs (Lonza, CA) were purchased at Passage 2 and cultured in mesenchymal stem cell basal medium (Lonza, CA) with mesenchymal cell growth supplement, L-glutamine, and GA-1000 (Lonza, CA), or DMEM, respectively and used up to Passage 7. Cell culture was performed in an incubator supplying 5% CO_2_ at 37 °C with medium changed every 3 days. For cell encapsulated hybprinting, cells were trypsinized and resuspended in the gelbrin bioink solution at 37 °C at a density of 2×10^6^ cells/mL before cooling. After printing and culturing, cell viability in gelbrin was evaluated by a fluorescent live-dead study at different time points. Briefly, at each time point, samples were washed with 1x PBS and stained by a mixture of calcein, AM (1 μL/mL; Invitrogen, MA) and ethidium homodimer-1 (2 μL/mL; Invitrogen, MA) for half an hour at 37 °C. The samples were then imaged under an inverted fluorescence microscope (BZ-X800, Keyence, Japan). Resulting images were then analyzed to quantify viability with ImageJ.

Cell proliferation was measured via metabolic activity using a 3-(4,5-dimethylthiazol-2-yl)-5-(3-carboxymethoxyphenyl)-2-(4-sulfophenyl)-2H-tetrazolium (MTS) assay kit (Abcam, UK). Briefly, the MTS reagent was added to each well containing one hybprinted sample at a ratio of 1:10 (MTS reagent:culture medium), and incubated for 2 hrs. 100 μL of the reagent and medium solution per sample was transferred to a 96 well-plate, and absorbance was quantified using a plate reader at OD 490 nm (SpectraMax iD3, Molecular Devices, CA).

### BSA patterning

For *in situ* crosslinking, FITC-BSA (Sigma, MO) was dissolved at 1% and mixed with 500 units/mL thrombin 1:1 to obtain a mixture of 0.5% FITC-BSA and 250 units/mL thrombin. For non *in situ* crosslinking, FITC-BSA was directly dissolved at 0.5%. The solutions were loaded onto a 384 well-plate for ADE deposition with a pattern of 6 parallel lines on half of a SE-printed gelbrin hydrogel (**Supplemental Figure S10d**). The droplet sizes were varied at 10, 50, and 100 nL per spot, with a total of 324 drops within the scaffold, correlating a total BSA amount of 16.2 81 and 162 μg, respectively. Upon crosslinking, hybprinted acellular samples were cultured in DMEM medium at room temperature, while cellular samples were cultured in DMEM medium in a 37 °C incubator.

### FITC-BSA release and retention

The amount of BSA released into PBS was quantified via fluorescence detection by collecting and replenishing the samples with fresh PBS at each time point. The fluorescence of the collected PBS was measured using a plate reader (SpectraMax iD3, Molecular Devices, CA) at excitation/emission wavelengths of 485/535 nm and compared against a standard curve.

At each time point, the same samples were also used to quantify BSA retention within the scaffold by direct fluorescence imaging. Briefly, after collecting PBS for the release study but prior to replenishing the samples with fresh PBS at each time point, the samples were imaged under an inverted fluorescence microscope (BZ-X800, Keyence, Japan). The fluorescence of the printed samples in different regions were quantified by integrating the gross fluorescent intensity in the region of interest using ImageJ.

### Hybprinting bone-tendon mimicking scaffold

A bone-tendon mimicking scaffold was fabricated by printing PCL as rigid “bony” part and gelbrin as soft “tendon” part following the previously mentioned hybprinting process. The PCL was printed into 2 separate 1 cm × 1 cm lattices, which were connected by a 1 cm × 6 cm hSMC-laden gelbrin lattics. The two parts had 0.5 cm overlap for soft-rigid integration. Printed scaffold had 5 layers with a thickness of 1 mm. 10 μg/mL FGF-2 was mixed with 500 units/mL thrombin 1:1 and patterned onto each layer of the gelbrin only part by the ADE module. The FGF-2/thrombin mix was patterned in a 6 by 36 array with 15 nL of solution on each spot per layer. The final amount of FGF-2 loaded in one scaffold was calculated to be 81 ng. Chondrogenic culture medium was prepared following previously described protocol, briefly consisting of DMEM, 100 nM Dexamethasone, 50 μg/mL Ascorbic-2-Phosphate, 1x Sodium Pyruvate, 1x Penicillin-Streptomycin, 5 μg/mL ITS premix and 10 ng/mL TGF-β3 ^118^. The hybrid scaffolds were cultured in chondrogenic culture medium for up to 7 days and analyzed for hMSC differentiation at each time point.

### RNA isolation and quantitative real-time PCR (qPCR)

qPCR analysis, following a protocol described in prior article ^119^, was used to evaluate hMSC differentiation into fibrochondrocytes. Briefly, at each time point, samples were collected and sectioned into PCL and gelbrin parts separately. Each sectioned sample was then treated with 1 mL of total RNA extraction reagent (R401-01, Vazyme, China). 1/5 volume of chloroform was added into the cell lysate, and the mixture was centrifuged at 12,000 g for 15 min at 4 °C. The upper aqueous phase was collected and mixed with an equal volume of isopropanol. The mixture was then centrifuged again at 12,000 g for 15 min at 4 °C. The precipitate was washed twice with 75% ethanol, air-dried, and resuspended in DNase/RNase-free water. RNA concentration and purity (A_260/280_ and A_260/230_) were measured using NanoDrop One (Thermal Fisher Scientific, MA). cDNA was synthesized from 1 μg RNA using a HiScript IV RT SuperMix kit (Vazyme, China) in a total volume of 20 μL in a thermal cycler (MiniAmp Plus, Applied Biosystems, MA). Reverse transcription conditions were as follows: 50 °C for 15 min followed by 85 °C for 5 s.

Expression of different genes were characterized with a quantitate real-time PCR machine (QuantStudio 6 Pro, Applied Biosystems, MA). A mixture of 0.4 μL cDNA, 0.5 μL forward primer, 0.5 μL reverse primer, 3.6 μL DNase/Rnase-free water and 5 μL Taq Pro Universal qPCR SYBR green master mix (Vazyme, China) was added into each reaction. PCR conditions were as follows: 95 °C for 5 min followed by 40 cycles of 95 °C for 15 s and 60 °C for 30 s. Primers used are listed in **Supplemental Table S13** with *ACTB* as the housekeeping gene. Values were normalized to *ACTB*, and data are shown as fold differences relative to the reference sample set (2^-ΔΔCT^).

### Mechanical bioreactor fabrication and mechanical stimuli

The bioreactor was constructed using laser cutting, CNC milling, and 3D printing of clear 5 mm acrylic, aluminum, and PETG thermoplastic. Two 3D-printed pulleys were created using a Prusa MK4 printer for the timing belt system, while all exterior housing for the bioreactor was laser cut. All parts interfacing with the sample were milled out of aluminum to be anodized to create a tarnish-free, sterilizable, and biocompatible surface. The bioreactor was powered via a linear actuator controlled by an Arduino controller; movement could be calibrated by the user by inputting maximum displacement and frequency through an LCD interface. An *in situ* battery system provided power for the circuitry, enabling device autonomy so that it could operate in an incubator without the need for a power cord. To sterilize the bioreactor, all non-electrical components were sprayed with 70% ethanol; then, the electronics and rubber timing belt were wrapped in foil before the whole system was left under UV in a sterile hood for 2 hours. Foil coverings were applied to preserve the integrity of the belt’s teeth during UV sterilization. After UV sterilization, the bioreactor was considered sterile. Samples could be loaded onto the bioreactor to undergo cyclic, gradual, or static loading. For slow stretch, samples would experience 1% stretch every 24 hours to mimic the bone-tendon development rate. Sample media was refreshed every 3 days. Once the experiment was complete, samples were unloaded from the bioreactor in a hood using sterile tweezers to be evaluated via RT-qPCR.

### Statistical analysis

Experiments are performed with at least three replicates unless otherwise noted. All statistical analyses were performed using Prism 7.0 (GraphPad Software Inc.). Quantitative data were presented as mean and standard error of mean (SEM). For qPCR study, multiple comparisons were analyzed with ANOVA test, with the level of significance was set at p < 0.05.

## Notes

### Competing Interest Statement

The authors have declared no competing interest.

