## Supplemental file for "Development of a Novel Hybprinter-SAM for Functionally Graded Tissue Engineering Constructs with Patterned and Localized Biochemical Signals"

**Supplementary Information**

Figure S1. Pictures showing process flow for Hybprinter-SAM.
Figure S2. Process flow of hybprinting design and fabrication.
Figure S3. Various demonstrations fabricated by Hybprinter-SAM.
Table S4. Important printing parameters for Hybprinter-SAM.
Figure S5. Crosslinking mechanism of gelbrin hydrogel network.
Figure S6. Difference during 14-day culturing between cellular and acellular scaffolds.
Figure S7. Mechanical measurement setup for the hybprinted scaffold.
Figure S8. COMSOL simulation of temperature change in the hybprinted scaffold.
Figure S9. MTS proliferation study for 1 mm thick samples.
Figure S10. Fluorescence retention of patterned FITC-BSA onto SE printed gelbrin scaffold.
Figure S11. Design and setup of custom designed mechanical bioreactor.
Table S12. Mechanical stimulus types and rates that can be provided by the custom designed bioreactor.
Table S13. Information of the primers used for qPCR.

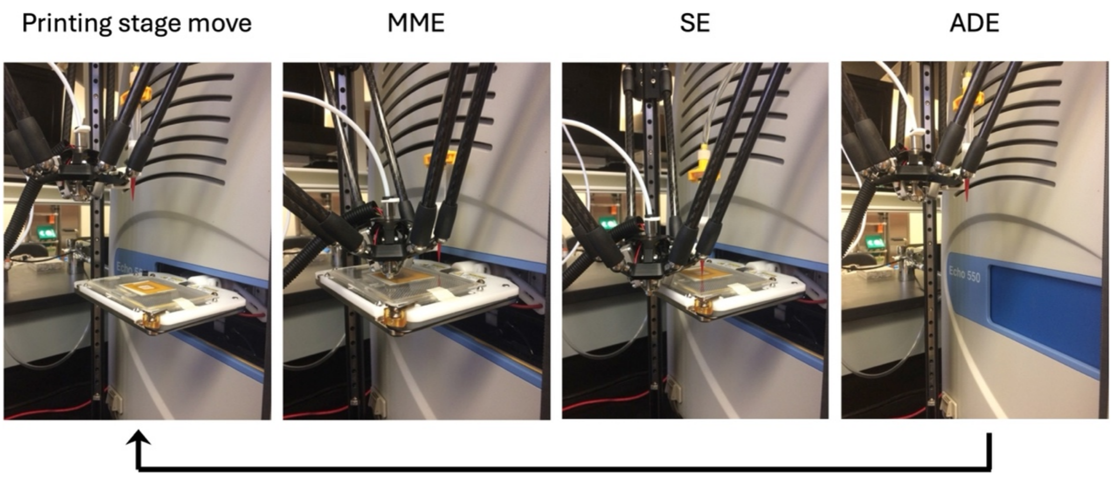


**Supplemental Figure S1.** Pictures showing process flow for Hybprinter-SAM, following a layer-by-layer fabrication protocol in the order of MME→SE→ADE.

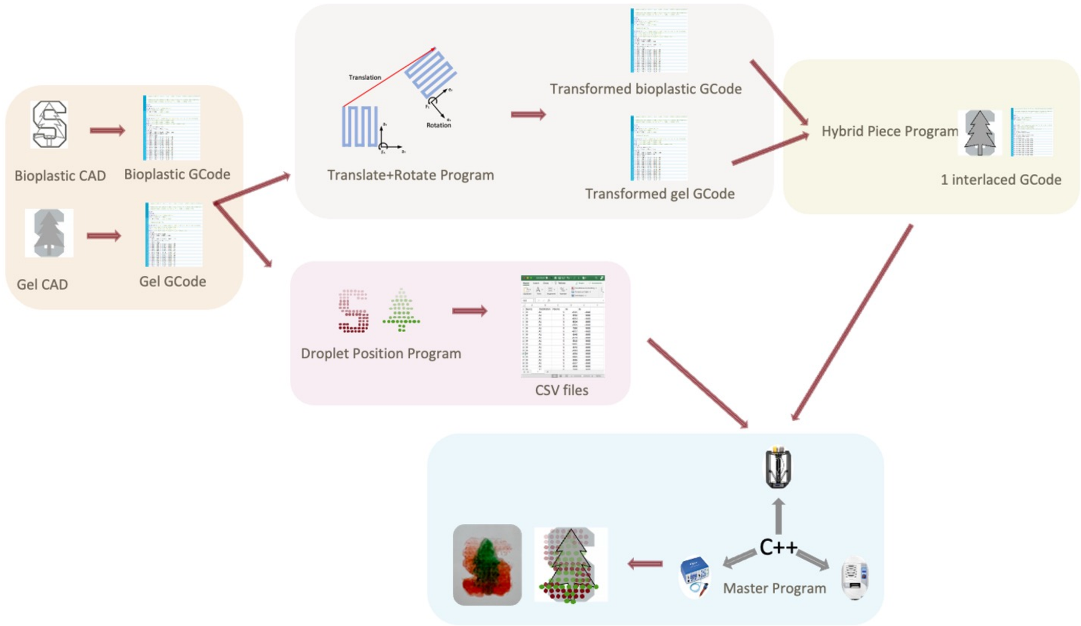


**Supplemental Figure S2.** Process flow of hybprinting design and fabrication including CAD design, slicing, coordinate system transformation, droplet pattern generation, and final master programming integration.

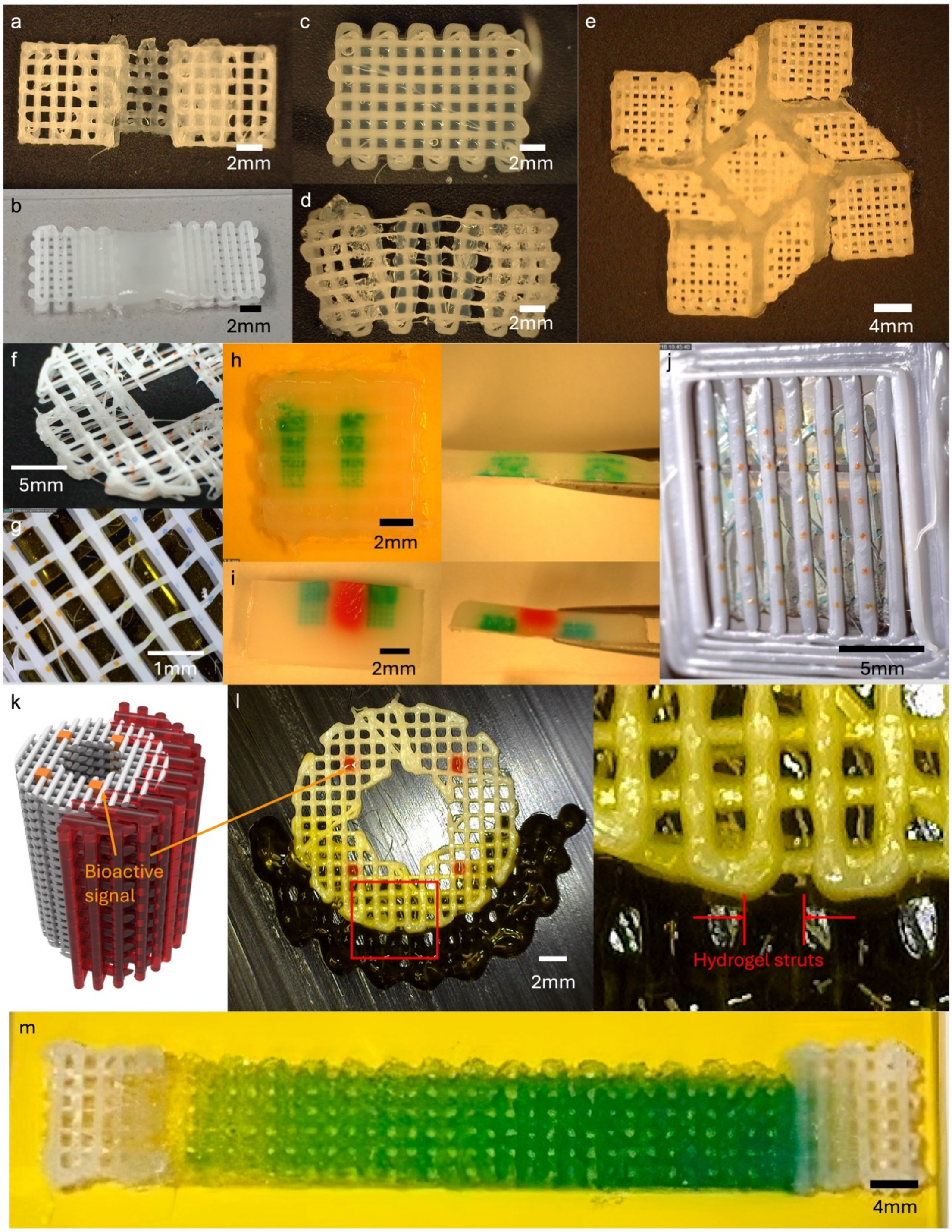


**Supplemental Figure S3.** (a-e) Hybrid soft-rigid tissue constructs of hydrogel and PCL that can withstand robust mechanical manipulation. (f, g) ADE deposition of droplets onto MME printed PCL struts. (h, i) Hydrogels with biological signal patterns. (j-l) Soft-rigid hybrid constructs with multiple biological factors (demonstrated by dyes) with different patterns, enabled by Hybprinter-SAM. (m) Printed sample of the soft-rigid construct with biological gradient, represented by color.

| Materials | Printing parameters | Values |
|---|---|---|
| PCL | Printing speed | 5 mm/s |
| | Printing temperature | 120 °C |
| | Extrusion rate | 100%-400% |
| | Layer height | 0.2 mm |
| Gelbrin | Printing speed | 2 mm/s |
| | Printing temperature | 20 °C |
| | Extrusion pressure | 10-20 psi |
| | Layer height | 0.2 mm |

**Supplemental Table S4.** Important printing parameters for Hybprinter-SAM. Note that extrusion rate for PCL was a parameter set for controlling the actual strut size printed with MME.

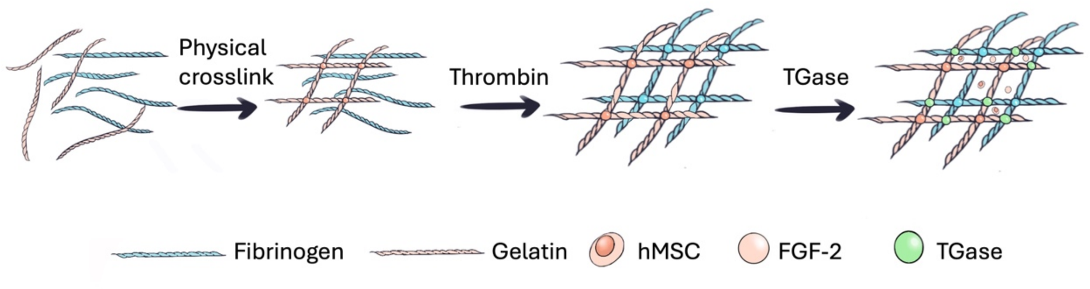


**Supplemental Figure S5.** Crosslinking mechanism of gelbrin hydrogel network: Gelatin first underwent temperature dependent physical crosslinking before printing; upon printing, thrombin first crosslinked fibrinogen into fibrin, followed by $Ca^{2+}$-dependent TGase crosslinking of gelatin and fibrin.

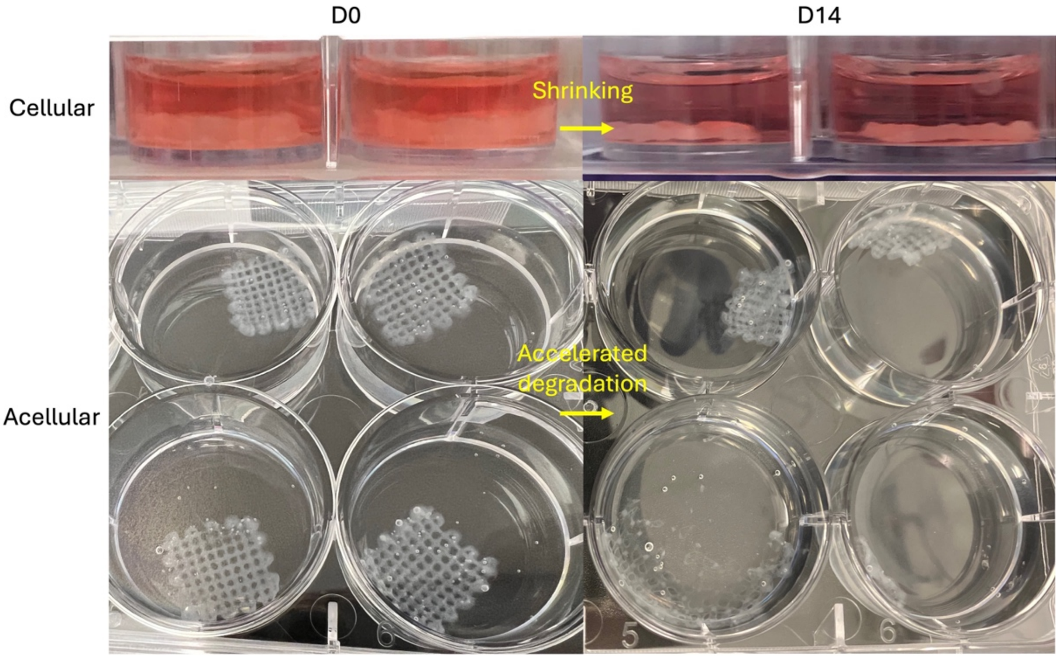


**Supplemental Figure S6.** Difference during 14-day culturing between cellular and acellular scaffolds, where cellular scaffolds shrank yet maintained its structural integrity, and acellular scaffolds almost fully degraded and dissembled.

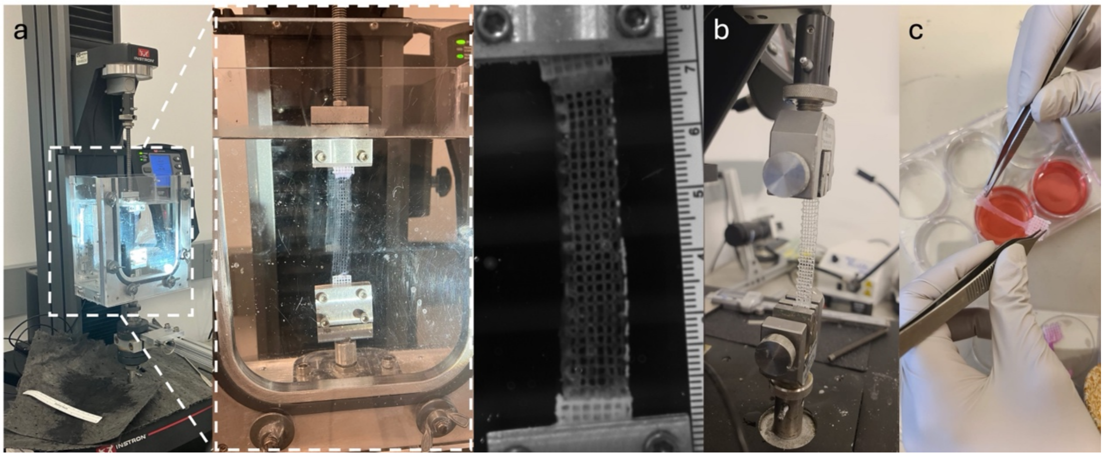


**Supplemental Figure S7.** Mechanical measurement setup for the hybprinted scaffold. (a) Measurement setup in water bath. (b) Measurement setup in air. (c) Gel deformation in air affecting measurement.

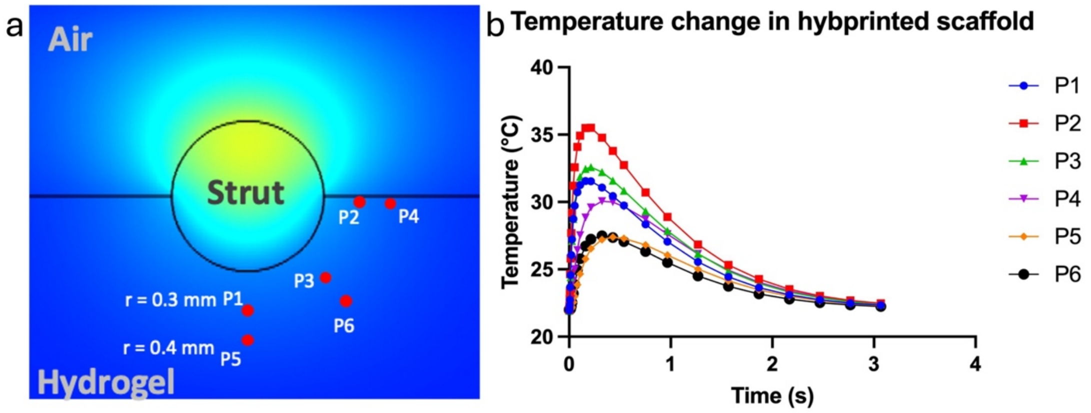


**Supplemental Figure S8.** COMSOL simulation of temperature change in the hybprinted scaffold. (a) Simulation design. (b) Simulation results for probes at different locations.

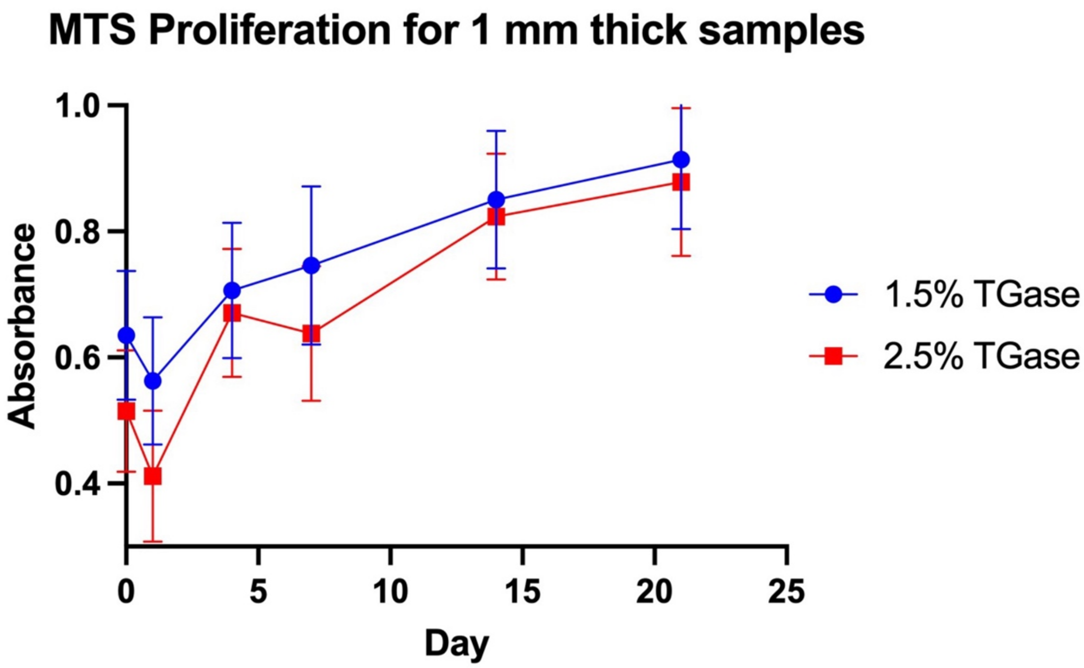


**Supplemental Figure S9.** MTS proliferation study for 1 mm thick samples.

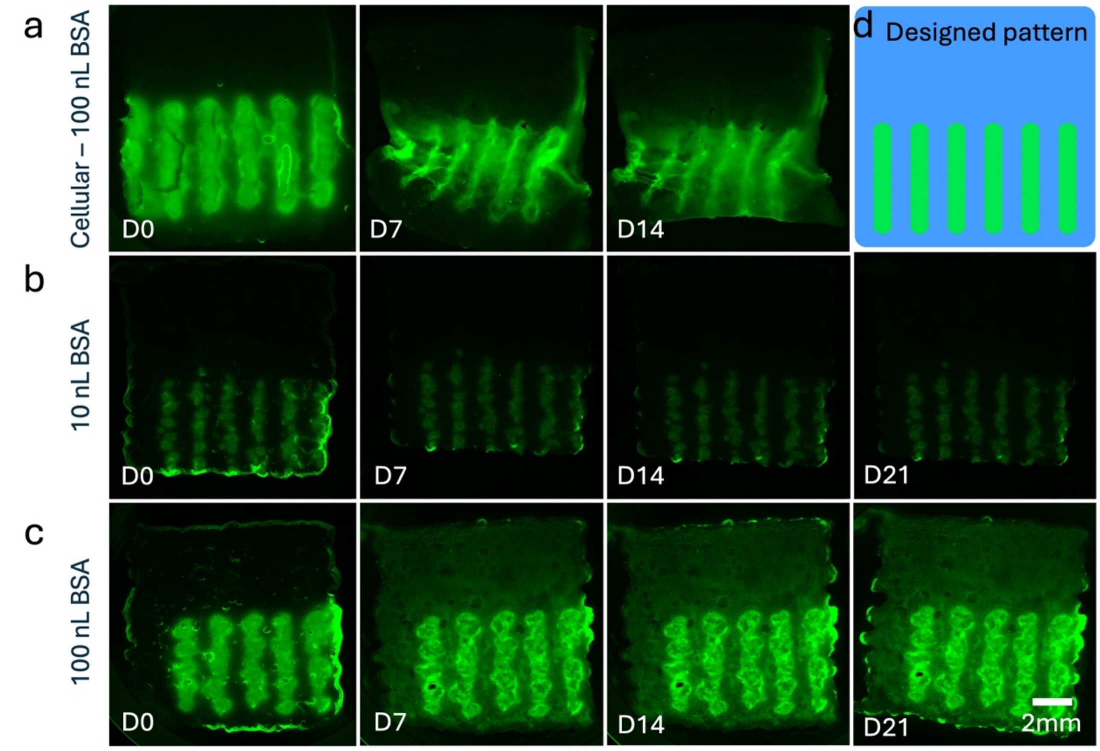


**Supplemental Figure S10.** (a-c) Fluorescence retention of patterned FITC-BSA onto SE printed gelbrin scaffold. (a) Cellular condition, 100 nL BSA droplet size. (b) Acellular condition, 10 nL BSA droplet size. (c) Acellular condition, 100 nL BSA droplet size. (d) Conceptual design of the pattern.

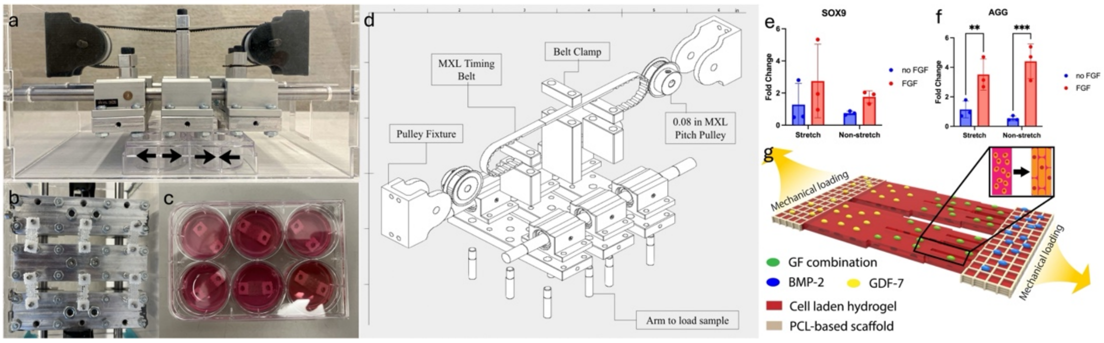


**Supplemental Figure S11.** (a) Side view of the bioreactor compatible with a 6-well plate. (b-c) Samples in a 6-well plate loaded onto the bioreactor. (d) An exploded schematic of the bioreactor's hardware and working mechanism. (e-f) qPCR expression of (e) SOX9 and (f) AGG, (e) COL1 and (f) AGG for comparison of mechanically stretched and non-stretched groups at D7. (g) Design for high-throughput screening comprising mechanical stimuli, soft-rigid material integration, combinatorial growth factors loading to study bone-tendon regeneration, where one may deposit BMP-2 for bone development, GDF-7 for tendon development, and screen different growth factors combinations for fibrocartilage development.

| Mechanical Stimuli Type | Physiological Strain/Rate | Relevant Frequency |
|---|---|---|
| Static Tension | 5% | NA |
| Slow Tension | 0.1 mm/day | NA |
| Cyclic Tension | 11% | 0.5 - 1 Hz |
| Cyclic Tension-Compression | 11% | 0.5 - 1 Hz |

**Supplemental Table S12.** Mechanical stimulus types and rates that can be provided by the custom designed bioreactor.

| Gene | FW | RV |
|---|---|---|
| ACTB | CACCATTGGCAATGAGCGGTTC | AGGTCTTTGCGGATGTCCACGT |
| SOX9 | GACTTCCGCGACGTGGAC | GTTGGGCGGCAGGTACTG |
| SCX | CCCAAACAGATCTGCACCTTC | GCGAATCGCTGTCTTTCTGTC |
| COL1 | GGACACAGAGGTTTCAGTGGT | GCACCATCATTTCCACGAGC |
| COL2 | TGGGGCCTTGTTCACCTTTGA | CGAGGCAACGATGGTCAGCC |
| AGG | TCGAGGACAGCGAGGCC | TCGAGGGTGTAGCGTGTAGAGA |

**Supplemental Table S13.** Information of the primers used for qPCR.
